# Kidzoi Enables Cell-Type-Specific Regulatory Activity and Variant Effect Prediction in Kidney

**DOI:** 10.64898/2026.09.28.746288

**Authors:** Arif A. Rather, Dongwon Lee

## Abstract

Sequence-to-function (S2F) deep learning models predict genome-wide functional profiles directly from DNA sequences and have significantly advanced our understanding of gene regulation. However, existing S2F models are fundamentally constrained by their initial training datasets, and no single model encompasses the full spectrum of cell types, clinical conditions, and cellular states, while training new models *de novo* is nontrivial. Here, we develop Kidzoi, a kidney cell-type-specific S2F model obtained by transfer learning from the pretrained model Borzoi using single-nucleus chromatin accessibility data from human kidney tissue. Kidzoi accurately predicts cell-type-specific chromatin accessibility and improves regulatory variant effect prediction in proximal tubule cells by up to 23% in terms of *AUROC* over existing models. Furthermore, Kidzoi effectively prioritizes genome-wide association study (GWAS) fine-mapped causal variants associated with kidney function (estimated glomerular filtration rate based on both creatinine and cystatin C; eGFRcr-cys) while resolving their cell-type specificity. Systematic ablation of flanking sequence indicates that a local context of 32–64*kbp*, representing only 6–12% of the full model input, captures most of the information needed for variant effect prediction. Finally, we show that a multi-task model achieves comparable performance to single-task models at a substantially reduced computational cost.

## 1 Introduction

Deep learning models trained on DNA sequences hold great promise for deciphering the grammar that governs cis-regulatory element activity. These sequence-to-function (S2F) models are trained on large, diverse experimental compendia spanning multiple modalities, such as gene expression (Cap Analysis of Gene Expression or CAGE, RNA-seq), chromatin accessibility, transcription factor binding, and histone modification, to predict these profiles directly from DNA sequences. A central application is variant effect prediction: by comparing predictions for reference and alternative alleles, these models score the functional consequences of non-coding variants [1–6], offering an efficient *in silico* means to prioritize disease-associated variants [7–13]. These models also enable *in silico* design of synthetic regulatory elements with desired activity [14–17]. This capability stems from their ability to learn hierarchical sequence representations from short transcription factor binding motifs to broader compositional patterns directly from raw DNA sequence without relying on otherwise handcrafted features [18,19].

Although modern S2F models represent many tissues, cell types, and assays, they capture only a subset of biological contexts. Consequently, investigators studying specialized tissues, cell types, or clinical conditions not represented in their training sets have to either settle for general-purpose models as a proxy, or train their own models *de novo*. However, training such models from scratch is not a trivial task as numerous design choices (e.g., input length, target resolution, architecture, initialization, optimization, and, critically, which data modalities to include) can strongly affect performance and generalization [20–25]. For example, while expression quantitative trait loci (eQTL) prediction benefits from multi-modal data training, chromatin accessibility variant prediction benefits most from models optimized on accessibility data alone [6]. Transfer learning offers an efficient alternative. Because pretrained S2F models already encode the regulatory grammar of the genome in their weights, they can be adapted to new cell types and conditions without training *de novo*, lowering cost while improving accuracy [26–29].

We developed a kidney cell-type-specific S2F model by fine-tuning the pretrained Borzoi model [5] on single-nucleus ATAC-seq (snATAC-seq) data from human kidney tissues [30]. To determine whether our transfer learning strategy generalizes beyond Borzoi, we repeated the same procedure using the pretrained Enformer model [1]. The resulting models are referred to as Kidzoi (Kidney-Borzoi) and Kidformer (Kidney-Enformer), respectively, owing to their derivation from the Borzoi and Enformer architectures and their specialization for modeling kidney cell-type-specific chromatin accessibility profiles. Briefly, Borzoi and Enformer address the same prediction task, but they differ in their training data, input sequence length, output resolution, and architecture. Both are pretrained on large functional datasets, including chromatin accessibility, transcription factor binding, histone modifications, and CAGE; Borzoi is additionally trained on RNA-seq data. Enformer uses an encoder–transformer architecture with a 196,608*bp* input and predicts the central 114,688*bp* at 128*bp* resolution. In contrast, Borzoi uses a U-Net–transformer architecture [31,32] with a 524,288*bp* input, models the central 196,608*bp*, and predicts at 32*bp* resolution.

We systematically evaluated Kidzoi and Kidformer models across three distinct tasks: (1) predicting ubiquitous and cell-type-specific accessibility across ten different human kidney cell types, (2) predicting regulatory variant effects measured by allelic imbalance in chromatin accessibility at heterozygous sites, and (3) prioritizing genome-wide association study (GWAS) fine-mapped variants associated with kidney function. We benchmarked our models against a kidney-specialized model (ChromKid) and general-purpose models (AlphaGenome, Borzoi, Enformer, Sei) [1,3,5,6,30]. We further asked how much sequence context is required by the models to achieve optimal performance, by ablating flanking sequence, and whether multi-task training sacrifices accuracy relative to single-task models.

## 2 Results

### 2.1 Kidzoi improves prediction of cell-type-specific accessibility

To build kidney-specific S2F models, we adapted pretrained Borzoi and Enformer models to human kidney snATAC-seq datasets using a transfer learning strategy (**Fig. 1A,B**). We replaced each pretrained model’s output head with a kidney-specific head predicting accessibility across ten kidney cell types. This involved adding a 1 × 1 convolution over Borzoi’s 1,920-dimensional representations and a linear layer over Enformer’s 3,072-dimensional embeddings, matching each backbone’s native design. As training targets, we used the snATAC-seq data available from Loeb et al. [30], the same dataset used to train the ChromKid model. To avoid data leakage, we strictly maintained the original splits (i.e., training, validation, and test sets) defined during the pretraining of Enformer and Borzoi. Because Borzoi and Enformer were not trained on identical sequence coordinates, the specific genomic regions included in each split differ between the two models, leading to differences in the sequences used for training, validation, and testing.

**Figure 1:**
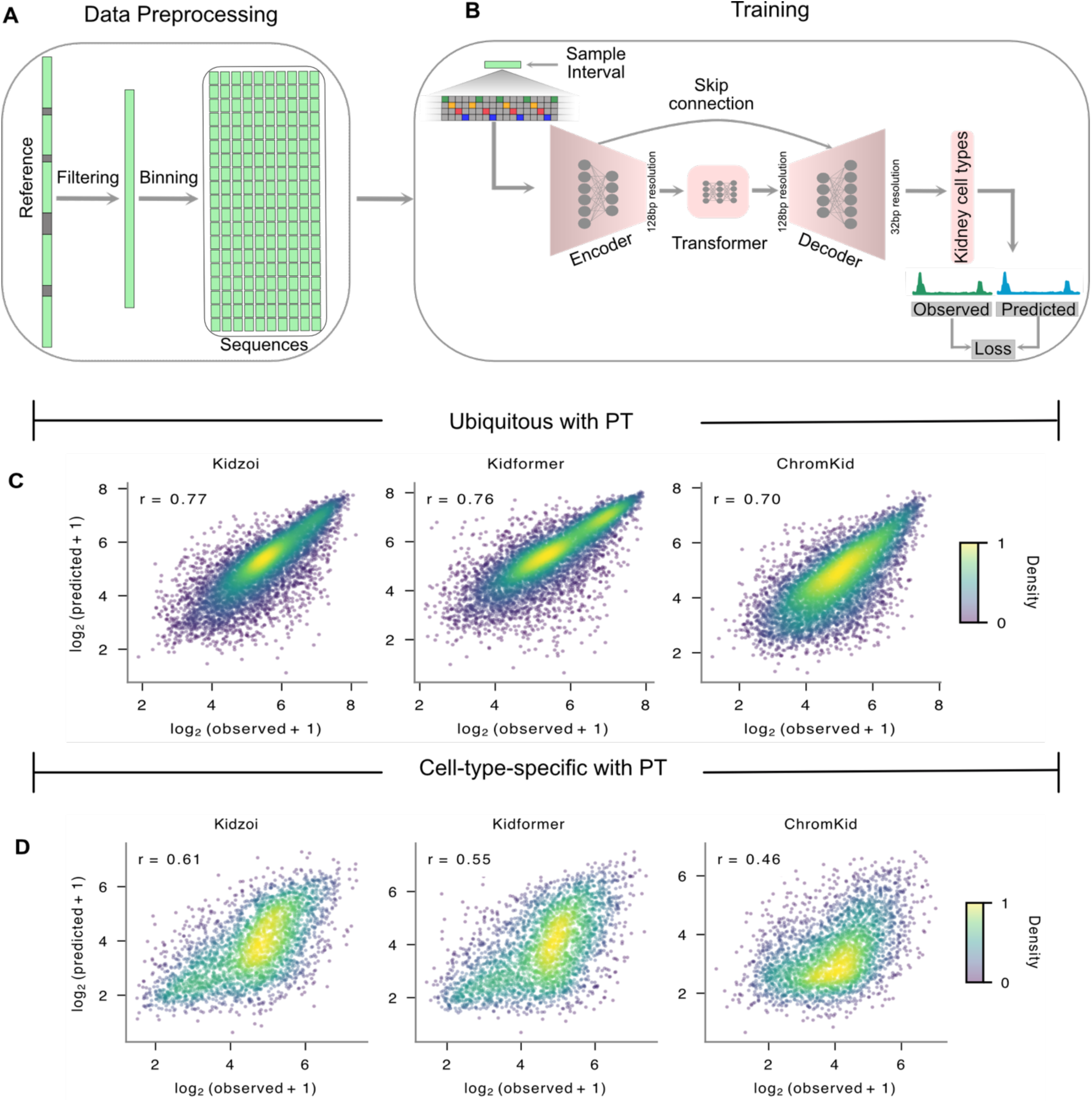
Model development and accessibility prediction. **(A)** Data preprocessing: the reference genome is filtered (assembly gaps, ENCODE blacklist), partitioned into fixed-length bins, and each bin is paired with its DNA sequence and matched bigWig signal. **(B)** Architecture and transfer strategy for Kidzoi: a U-Net encoder-decoder with transformer layers compresses the input to 128*bp*, captures long-range interactions, and reconstructs predictions at 32*bp*; a kidney-specific head outputs ten cell types. Shift and reverse-complement augmentation are applied during training. **(C)** Observed *vs*. predicted coverage in proximal tubule for ubiquitous test regions (Kidzoi, Kidformer, ChromKid). **(D)** As in C, for proximal-tubule-specific regions; performance drops across all models, with Kidzoi achieving the highest performance (*r* = 0.61).

We first quantified how effectively Kidzoi and Kidformer generalize to unseen genomic regions in the test set. To this end, we divided the kidney open chromatin regions in the test set into two groups: ubiquitous regions (accessible across all ten cell types) and cell-type-specific regions (accessible in one cell type and not in the rest), sourced from Loeb et al. [30]. The number of cell-type-specific peaks in the test set for Kidzoi ranges from 4,331 in stromal cells (Str) to 7,133 in distal tubule (DT), and the number of ubiquitous peaks is 8,450. For comparison, we evaluated Kidzoi and Kidformer models against the kidney-specialized (ChromKid) model [30]. In proximal tubule (PT), the most abundant cell type in the kidney, both Kidzoi and Kidformer predicted accessibility in ubiquitous regions with high accuracy (Pearson *r* = 0.77 and 0.76, respectively) and outperformed ChromKid (*r* = 0.70) (**Fig. 1C**).

Predicting cell-type-specific open chromatin regions is a substantially more challenging task as these regions are highly dependent on cell-state-specific regulatory programs and often involve complex combinations of transcription factor binding and distal regulatory interactions [33–35]. Nonetheless, Kidzoi outperformed Kidformer, achieving a relative gain of 11% in terms of Pearson *r* for PT-specific peaks (**Fig. 1D**; **Methods**). On ubiquitous peaks, the two models performed similarly in all ten cell types, and both outperformed ChromKid. Both models reached *r =* 0.67–0.84 *vs*. 0.56–0.73 for ChromKid for ubiquitous regions (**Fig. 2A**). Kidzoi outperformed Kidformer in predicting cell-type-specific peaks in every cell type except immune cells (Imm), achieving relative gains in Pearson *r* ranging from 6% (endothelial cells, End, and parietal epithelial cells, PE) to 59% (collecting duct, CD). Kidzoi also outperformed ChromKid, yielding relative gains of ∼10–68% (e.g., PE, *r* = 0.37 *vs.* 0.22) (**Fig. 2B**). The performance advantage of Kidzoi over Kidformer likely reflects its finer 32*bp* bin resolution, compared with Kidformer’s 128*bp*, which yields more fine-grained predictions. ChromKid generally yielded the lowest performance in predicting cell-type-specific regions, except in CD and loop of Henle (LOH), where it outperformed Kidformer.

**Figure 2:**
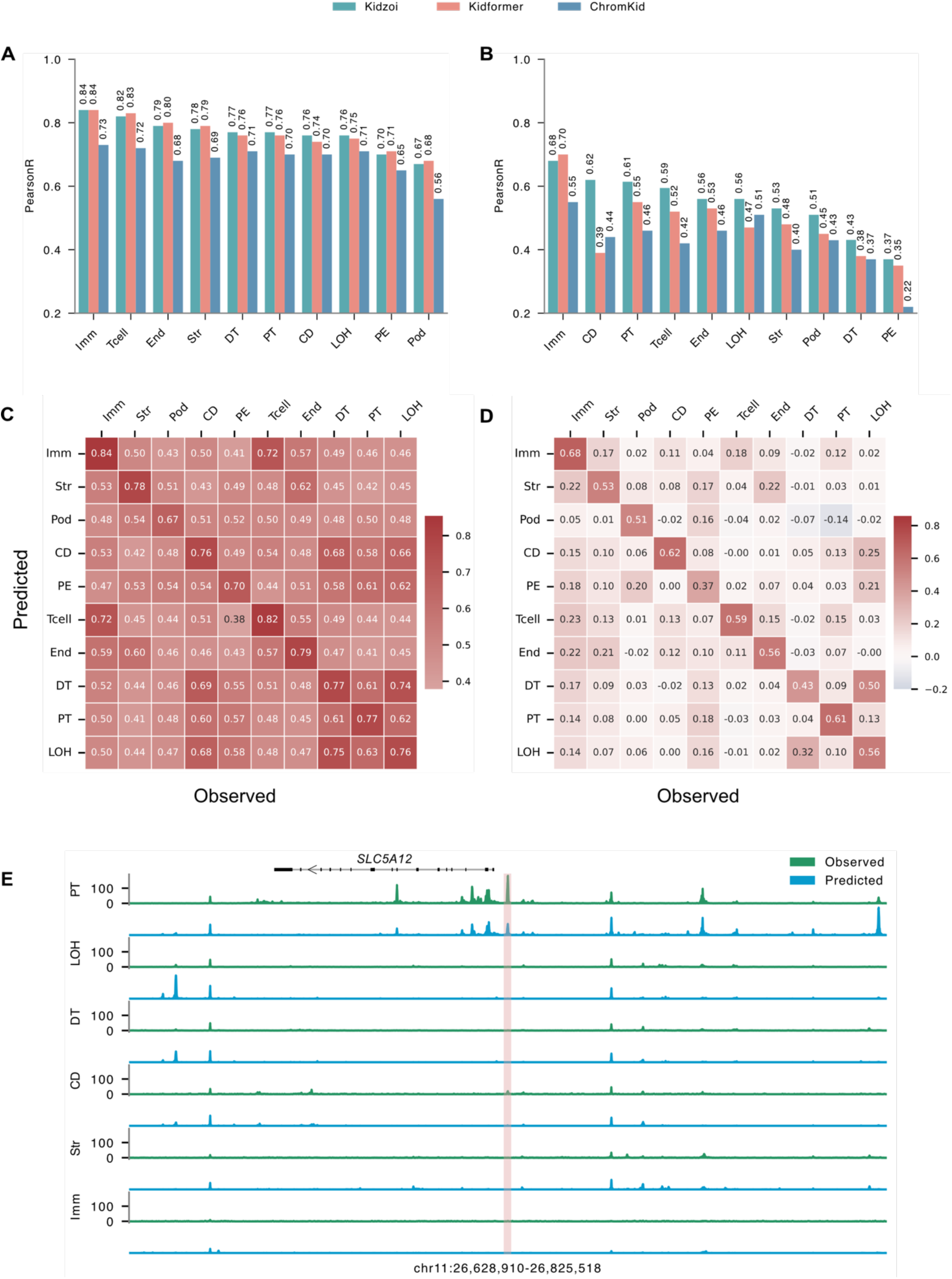
Accessibility prediction across cell types. **(A)** Ubiquitous-region performance across cell types. **(B)** Cell-type-specific-region performance. **(C)** Kidzoi cross-cell-type performance in ubiquitous regions: entry (*i, j*) is Pearson *r* between predicted signal in cell type *i* and the observed signal for cell type *j*; diagonal entries are within-cell-type, off-diagonal entries cross-cell-type. **(D)** As in C, for cell-type-specific regions. **(E)** Observed *vs*. predicted tracks across chr11:26,628,910–26,825,518 (hg38) near *SLC5A12* in six kidney cell types (Kidzoi). The shaded band marks the PT-specific region near *SLC5A12* described in the text. Cell-type abbreviations: Imm, immune; Str, stromal; Pod, podocyte; CD, collecting duct; PE, parietal epithelial; Tcell, T cell; End, endothelial; DT, distal tubule; PT, proximal tubule; LOH, loop of Henle. The same abbreviations are used for other figures.

We also assessed cell-type specificity of Kidzoi and Kidformer by evaluating each cell type’s prediction on every cell type’s observed chromatin accessibility. The results are summarized as 10 × 10 matrices for both ubiquitous regions and cell-type-specific regions (**Fig. 2C,D; Supplementary Fig. S1A,B**). Each entry (*i, j*) reflects the Pearson correlation between the predicted signal in cell type *i* and the observed signal in cell type *j*. Thus, diagonal entries represent prediction performance for matched cell types, while off-diagonal entries represent cross-cell-type prediction accuracy. When predicting ubiquitous regions, correlations were high both on and off the diagonal, as expected. However, we still achieved the highest correlation for the matched cell types (0.67–0.84), suggesting that even ubiquitous regions retain cell-type-specific signals. For cell-type-specific regions, however, high correlations were confined to the diagonal, except between the related DT and LOH cell types, with other off-diagonal correlations being mostly near 0, demonstrating that Kidzoi learned distinct regulatory signals per cell type. ChromKid, in contrast, showed elevated off-diagonal correlations, indicating that its predictions are less cell-type-specific (**Supplementary Fig. S1C,D**). As an illustration, Kidzoi correctly predicted a PT-specific region near *SLC5A12*, which encodes the sodium-coupled monocarboxylate transporter SMCT2 [36–38] (**Fig. 2E**).

### 2.2 Kidzoi improves regulatory variant effect prediction

S2F models can be used to score variants without additional fine-tuning by comparing predictions for reference and alternative alleles. We next tested whether the enhanced prediction accuracy in open chromatin regions of Kidzoi and Kidformer translates into improved variant effect prediction (VEP). We evaluated the performance of predicting allelic imbalance in chromatin accessibility between alleles at heterozygous sites in three major kidney cell types: PT, LOH, and DT. The data for this analysis were obtained from Loeb et al. [30]. The number of variants exhibiting allelic imbalance varied across cell types, ranging from 312 in DT to 2,417 in PT, with 1,360 in LOH. For each cell type, negative variants were sampled at a 7:1 ratio relative to the positive variants (see **Methods** for more details). As baseline models, we evaluated four general-purpose models (AlphaGenome, Borzoi, Enformer, and Sei) and a kidney-specific model, ChromKid. We scored each variant by averaging predictions over forward and reverse-complement sequences and over ±1*bp* shifts, and aggregated the signal in a 256*bp* window centered on the variant, corresponding to eight Kidzoi bins and two Kidformer bins (**Methods**). This 256*bp* aggregation window was maintained in all baseline models when possible. Our choice of this small window size is based on the highly localized cis-regulatory signal represented by allelic imbalance, which we examine below.

For each of the general-purpose models, we selected the top-performing track among the ATAC-seq and DNase-seq tracks, ranked by area under the receiver operating characteristic curve (*AUROC*). For Kidzoi, Kidformer, and ChromKid, we used the matched cell-type track. In PT, Kidzoi and Kidformer achieved *AUROC =* 0.82, which translates into relative gains of 23% over Enformer, Borzoi, and Sei (*AUROC* = 0.76), 10% over AlphaGenome (*AUROC* = 0.79), and 14% over ChromKid (*AUROC* = 0.78) (random baseline, *AUROC* = 0.5) (**Fig. 3A,C**). For each general-purpose model, we evaluated the area under the precision-recall curve (*AUPRC*) on the same track that had been selected as its top-performing track by *AUROC*. Both Kidzoi and Kidformer achieved *AUPRC* = 0.57, which translates into 29–56% relative improvement over the general-purpose models (*AUPRC*: 0.41–0.47), and 22% over ChromKid (*AUPRC* = 0.49) (random baseline = positive-class prevalence, 0.125) (**Fig. 3B,D**). DT and LOH showed a similar trend, with Kidzoi generally outperforming Kidformer, although in DT the best AlphaGenome track (*AUROC* = 0.83) scored slightly above Kidformer (0.82) (**Fig. 3C,D; Supplementary Fig. S2A–D**). Kidzoi and Kidformer generally outperformed ChromKid in effect-direction concordance across DT, PT, and LOH, based on both *AUROC* and *AUPRC*. Within this comparison, Kidzoi also outperformed Kidformer in *AUPRC* for DT and LOH (**Supplementary Fig. S2E,F**). For all cell types (PT, DT, and LOH), the highest-performing tracks in AlphaGenome, Enformer, and Borzoi corresponded to renal cell carcinoma, whereas the highest-performing Sei tracks were fetal kidney for PT and LOH and pancreas for DT. We next evaluated the cell-type specificity of Kidzoi by scoring allelic imbalance variants in PT using each of the ten kidney cell-type tracks. Kidzoi performed best in PT (**Fig. 3E,F**), matching the cell-type specificity of the variant effects. Moreover, Kidformer and ChromKid showed similar patterns (**Supplementary Fig. S3**). This result further suggests that the Kidzoi and Kidformer models effectively learn cell-type-specific regulatory features.

**Figure 3:**
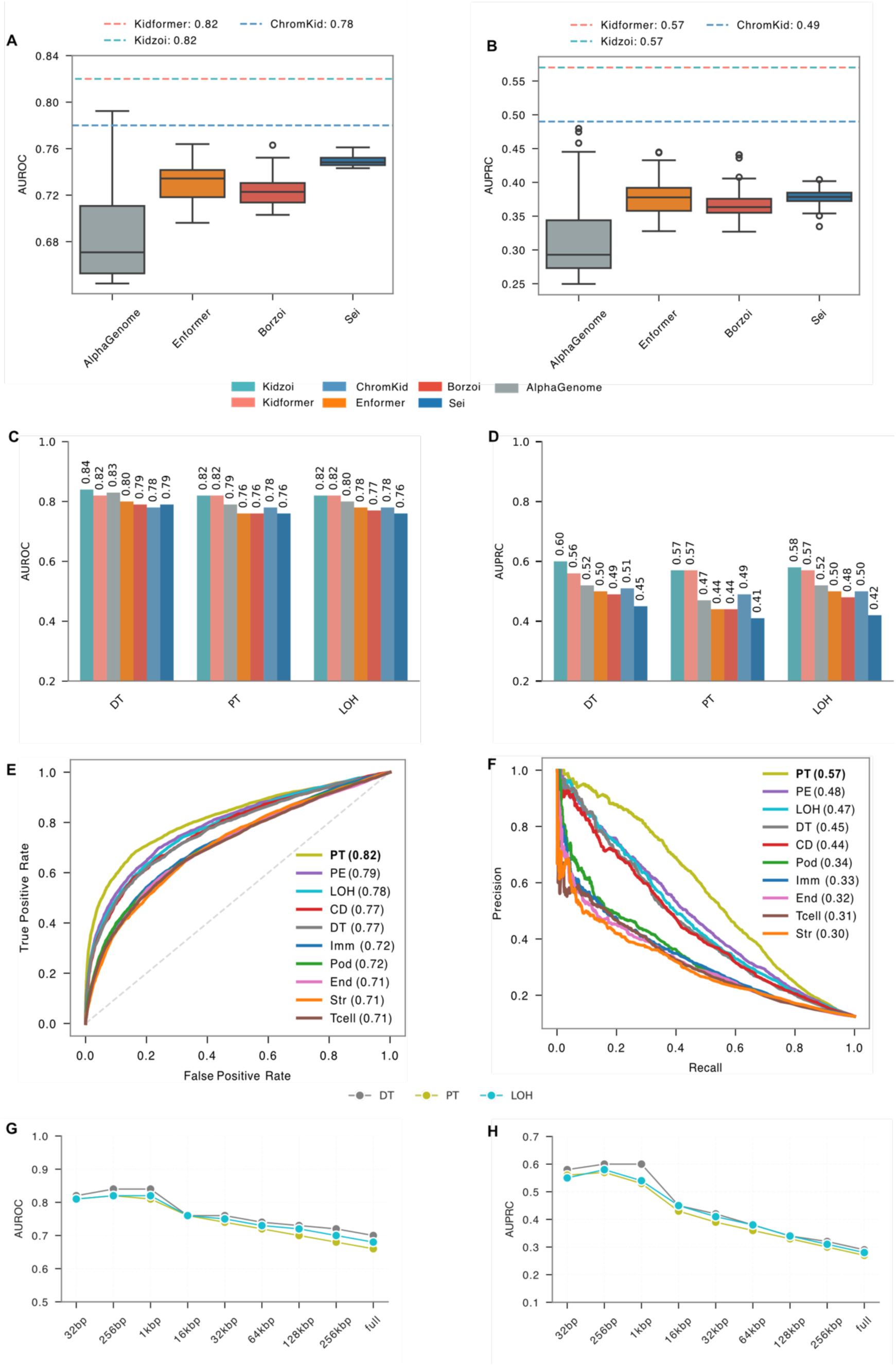
The performance of pretrained baseline models, ChromKid, Kidzoi, and Kidformer in predicting allelic imbalance in three major kidney cell types. Box plots (A) and (B) show performance distributions in predicting allelic imbalance in PT across the top 100 DNase and ATAC tracks, ranked by *AUROC*, from each pretrained model, while dashed horizontal lines indicate the performance of Kidzoi, Kidformer, and ChromKid as single reference values. (C, D) Bar plots showing the performance of AlphaGenome, Borzoi, Enformer, Sei, Kidzoi, Kidformer, and ChromKid in predicting allelic imbalance in terms of *AUROC* and *AUPRC* across DT, PT, and LOH. (E, F) ROC and precision-recall curves, respectively, for Kidzoi scoring of regulatory variants that show allelic imbalance in PT. (G, H) Line plots of Kidzoi *AUROC* and *AUPRC*, respectively, in predicting allelic imbalance as a function of signal aggregation window size around the variant position.

Lastly, we asked how model performance varies as a function of genomic window size for signal aggregation. For this, we considered the following window sizes centered on the variants: 32*bp*, 256*bp*, 1*kbp*, 16*kbp*, 32*kbp*, 64*kbp*, 128*kbp*, 256*kbp*, and the full sequence length, and computed *AUROC* and *AUPRC* at each. The model input sequence remained fixed at 524*kbp*; we only varied the output window used to aggregate the predicted signal around each variant. As shown in **Fig. 3G,H**, we achieved the best performance at window sizes between 256*bp* and 1*kbp*, after which performance decreased monotonically with increasing size across all three cell types (PT, DT, and LOH). This result indicates that predicted signal more than ∼0.5*kbp* from the variant adds little to the allelic imbalance prediction and that incorporating predicted distal genomic signals introduces noise that dilutes the local variant signal.

### 2.3 Kidzoi captures cell-type-specific effects of GWAS fine-mapped variants

Statistical fine-mapping of GWAS data can narrow down disease associations to smaller credible variant sets, but tight linkage disequilibrium (LD) blocks often mask true causal variants, and experimental screening at scale remains challenging [39–42]. S2F models that can predict cell-type-specific regulatory variants can be useful in prioritizing causal variant candidates from nearby variants in high LD, as these models rely only on sequence features. We thus asked whether Kidzoi and Kidformer models could discriminate non-coding fine-mapped variants associated with estimated glomerular filtration rate based on both creatinine and cystatin C (eGFRcr-cys), and, more importantly, whether they could uncover the specific cell types driving these GWAS signals. We used functionally informed fine-mapped variants from eGFRcr-cys GWAS, obtained from Loeb et al. [30]. After removing coding variants, we used 593 non-coding fine-mapped single nucleotide variants (SNVs) (posterior inclusion probability, PIP > 0.3) as positives and sampled minor allele frequency (MAF)-matched negatives at a 7:1 ratio from the same summary statistics with *P* > 0.5, excluding any variant within a fine-mapped locus (**Methods**). Using the LOH track, Kidzoi (*AUROC* = 0.79) and Kidformer (*AUROC* = 0.79) outperformed the matched kidney tracks of Borzoi (*AUROC* = 0.78, track: DNase, right kidney, male embryo) by 4%, Enformer (*AUROC* = 0.76, track: DNase, right kidney, male embryo) by 12%, Sei (*AUROC* = 0.78, track: DNase, fetal kidney) by 4%, AlphaGenome (*AUROC* = 0.76, track: DNase, right kidney) by 12%, and ChromKid (*AUROC* = 0.73, track: LOH) by 26% (**Fig. 4A**; **Supplementary Fig. S4A,C)**. Notably, none of AlphaGenome’s five highest-performing tracks corresponded to kidney tissue. On *AUPRC* (using the same matched kidney tracks selected for *AUROC* evaluation), Kidzoi and Kidformer models (0.46–0.48) were comparable to the matched tracks of AlphaGenome (*AUPRC* = 0.46), Borzoi (*AUPRC* = 0.47), and Enformer (*AUPRC* = 0.48) and exceeded ChromKid (0.39) and Sei (0.45) (**Fig. 4B**).

**Figure 4:**
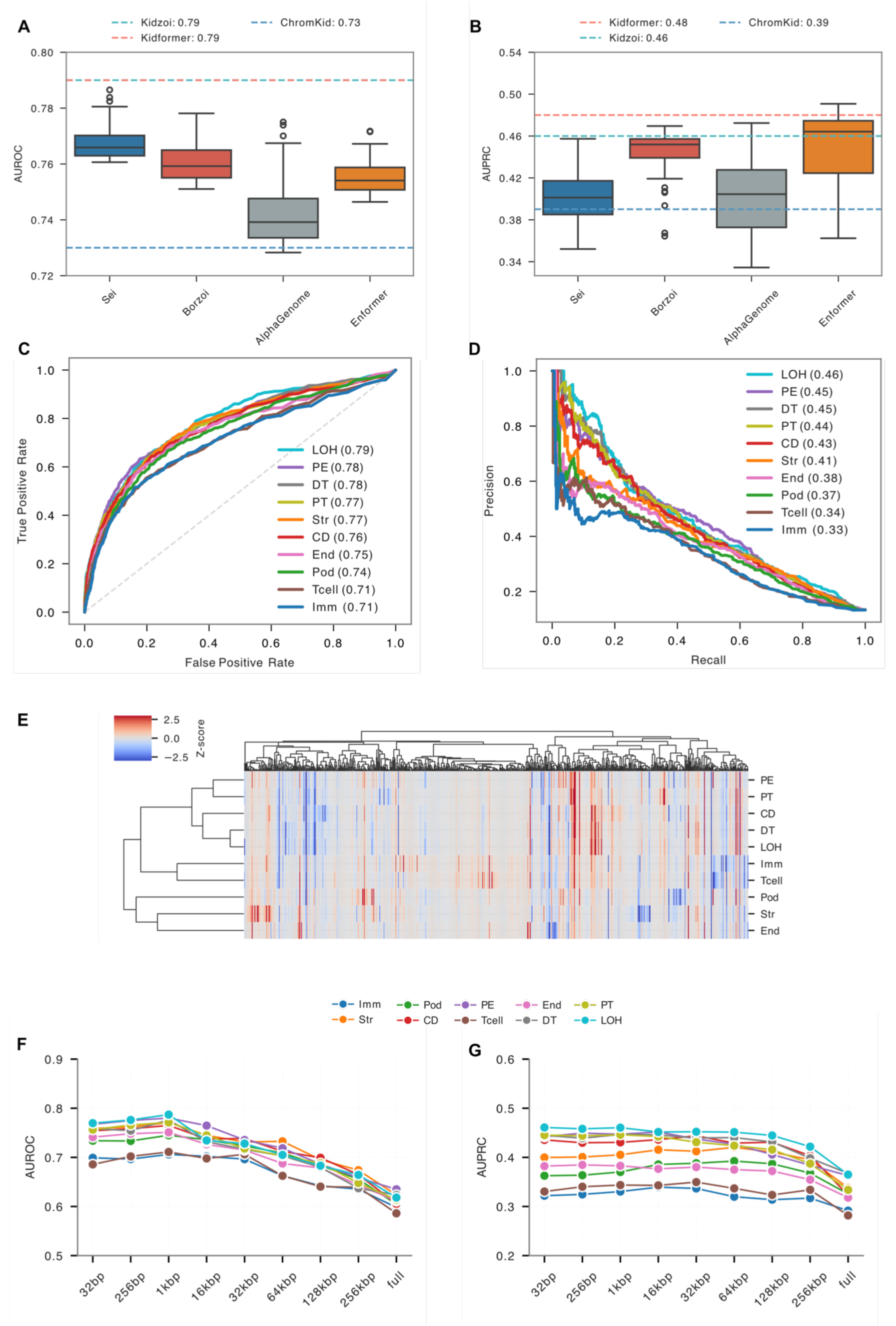
Performance on GWAS fine-mapped variants. **(A, B)** Box plots represent the performance on the top 100 DNase and ATAC tracks in terms of *AUROC* and *AUPRC*, respectively, while dashed horizontal lines indicate the performance of Kidzoi, Kidformer, and ChromKid as single reference values. **(C, D)** ROC and precision-recall curves for GWAS fine-mapped variants by Kidzoi across all ten cell types. **(E)** A heatmap of predicted effects of fine-mapped variants (columns) across the ten cell types (rows). Allelic log_2_ fold changes were computed as log_2_(ALT + 1) − log_2_(REF + 1) from cell-type-specific predicted chromatin accessibility scores, then z-scored within each cell type. Red indicates a predicted gain of accessibility in the alternative allele relative to that cell type’s average effect; blue indicates a relative loss. Hierarchical clustering using correlation distance was applied to both variants and cell types. **(F, G)** Line plots showing the *AUROC* and *AUPRC*, respectively, of Kidzoi as a function of signal aggregation window size around the variant position across all ten cell types.

This precision ceiling highlights important characteristics of complex trait genetics. Two factors can confound discriminative performance in this task. First, fine-mapped variants are not a perfect positive set; statistical fine-mapping yields credible sets that still contain non-causal variants, and even causal variants need not act through chromatin-accessibility regulation (some affect splicing, transcript stability, or other modalities) [41,43]. Second, complex traits reflect physiological contributions spread across diverse tissues and cell types [44], while our LOH model focuses on one specific kidney cell type, capturing only the LOH-attributable portion of the trait variance. For these reasons, no single cell-type model can be expected to rank an entire credible set above background, and not achieving the highest precision is fully consistent with the cell-type-specific architecture of regulatory variation.

The unique value of our Kidzoi and Kidformer models thus lies in their ability to resolve the precise cell types in which fine-mapped variants exert their function. Kidzoi achieved generally high performance in tubular epithelial cells (*AUROC* = 0.76–0.79; *AUPRC* = 0.43–0.46) (**Fig. 4C,D**), consistent with the established enrichment of kidney-function heritability in tubular epithelial candidate cis-regulatory elements [30]. Kidformer performed comparably, with slightly higher *AUPRC* in every cell type (**Supplementary Fig. S4A,B**). Computing allelic log_2_ fold changes for every fine-mapped variant across all ten cell types and performing unsupervised hierarchical clustering on the predicted effects alone grouped tubular and parietal epithelial cells (PT, CD, DT, LOH, PE) into one cluster, joined next by immune and T cells, with podocytes, stromal, and endothelial cells forming a separate cluster (**Fig. 4E**). These effects were sparse and cell-type-restricted rather than uniform, indicating that individual fine-mapped variants exert function within specific kidney compartments and cell types. Using the 90^th^ percentile cutoff of the distribution of absolute log_2_ fold changes pooled across all fine-mapped variants and cell types, approximately 30% of such variants are predicted to have a large impact in at least one kidney cell type. These variants, on average, are predicted to impact only three out of the ten distinct cell types. In summary, these results suggest that Kidzoi and Kidformer models partition GWAS signal across kidney cell types.

Lastly, we repeated the genomic aggregation window analysis using fine-mapped GWAS variants, following the same procedure used for allelic-imbalance variants. As shown in **Fig. 4F** and **Supplementary Fig. S4E**, *AUROC* peaked at or near the 1*kbp* window (32 bins in Kidzoi and 8 bins in Kidformer) and declined as the window size increased. *AUPRC* was less sensitive to window size (**Fig. 4G; Supplementary Fig. S4F**). Beyond 1*kbp*, it did not improve in PT, DT, LOH, PE, or endothelial cells with Kidzoi, or in any cell type with Kidformer, whereas Kidzoi *AUPRC* rose slightly at 16–64*kbp* in stromal cells, podocytes, CD, immune cells, and T cells before declining at larger windows. This result suggests that aggregating signals over larger genomic regions does not generally improve the identification of GWAS functional non-coding variants as it dilutes their local regulatory effects, consistent with the allelic-imbalance variant prediction.

### 2.4 Local sequence context is sufficient for variant effect prediction

Since the informative predictive signal for VEP is local **(Fig. 3G,H)**, we next asked how much flanking input sequence the models actually need for best performance. In other words, although Kidzoi accepts 524,288*bp* of input, it is unclear how much of this context contributes to prediction. We therefore ablated flanking sequence by retaining a centered window of increasing length and masking the remainder with *Ns*, and measured performance across three prediction tasks: open chromatin regions, allelic imbalance in chromatin accessibility, and fine-mapped GWAS variants (**Fig. 5**). Unlike varying the output window over which predictions are aggregated, this directly probes the input context the models need for prediction. To obtain predictions, each ubiquitous region was evaluated with a single forward pass, whereas cell-type-specific regions were evaluated separately for each cell type, as the sets of active regions differ across cell types. For VEP, each variant was evaluated using both the forward and reverse-complement sequences, and the resulting predictions were averaged; the ±1*bp* shift averaging used in the main analyses was omitted here to limit computational cost, so absolute values differ from those in **Figs. 3 and 4**.

**Figure 5:**
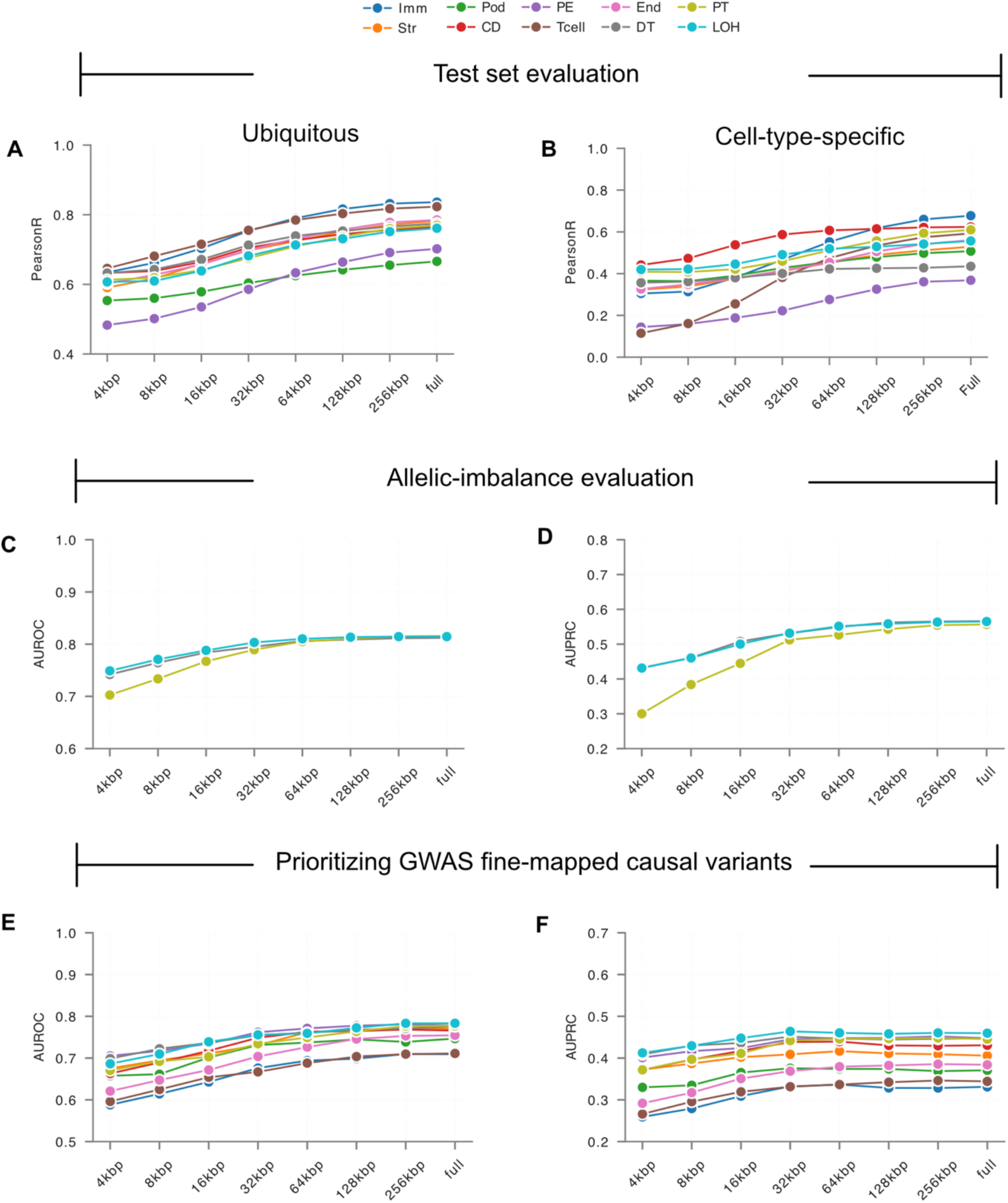
The effect of input sequence truncation on Kidzoi performance. The x-axis shows the length of the actual genomic sequence around the region or variant, with the remaining positions padded with *Ns* to maintain the full 524*kbp* input length. This allows assessment of model performance as a function of sequence context length using different metrics. **(A, B)** Performance in predicting ubiquitous and cell-type-specific regions, respectively. **(C, D)** *AUROC* and *AUPRC* for predicting allelic imbalance. **(E, F)** *AUROC* and *AUPRC* for predicting GWAS fine-mapped variants.

For open chromatin region prediction, most gains were achieved by expanding input context to 128*kbp*. Ubiquitous regions showed improvements in Pearson correlation from 0.48–0.65 at 4*kbp* to 0.64–0.82 at 128*kbp*, whereas cell-type-specific regions improved from 0.12–0.44 to 0.33–0.62 across cell types (**Fig. 5A,B**). Further increasing context from 128*kbp* to 524*kbp* yielded diminishing returns, with relative improvements of only ∼2–6% for ubiquitous regions and ∼2–13% for cell-type-specific regions (**Fig. 5A,B**; **Supplementary Tables S1 and S2**). For allelic imbalance prediction, most performance gains were achieved by expanding context from 4*kbp* to 32*kbp*. *AUROC* increased from 0.70–0.75 at 4*kbp* to 0.79–0.81 at 32*kbp*, with smaller further gains at the full 524*kbp* context (0.81–0.82; up to 9% relative gain). Similarly, *AUPRC* increased from 0.30–0.43 at 4*kbp* to 0.51–0.54 at 32*kbp* and showed only modest gains at 524*kbp* (0.56) across DT, PT, and LOH cell types. These results indicate that allelic imbalance prediction largely saturates at ∼32*kbp* of sequence context (**Fig. 5C,D**; **Supplementary Tables S3 and S4**). For GWAS fine-mapped variant prioritization, increasing sequence context substantially improved performance from 4*kbp* to intermediate context lengths. *AUROC* increased from 0.59–0.71 at 4*kbp* to 0.69–0.77 at 64*kbp* across the ten cell types, with further expansion to 524*kbp* providing additional but smaller gains (0.71–0.78). In contrast, *AUPRC* largely saturated by 32*kbp* (0.33–0.46), with minimal additional benefit at the full 524*kbp* (−2% to 6% relative change). These results indicate that most of the regulatory information required for GWAS variant prioritization is captured within tens of kilobases of sequence context (**Fig. 5E,F**; **Supplementary Tables S5 and S6)**. For allelic imbalance and for GWAS *AUPRC*, 32–64*kbp* (only 6–12% of the 524*kbp* input) captured most of the predictive signal; GWAS *AUROC* continued to improve modestly up to the full input. Longer context helps slightly, but most of the sequence context needed to predict regulatory variant effects lies within 64*kbp*. Kidformer behaved broadly similarly, although its allelic-imbalance *AUROC* was already high at 4*kbp* (**Supplementary Fig. S5**).

### 2.5 Single-task and multi-task model evaluation

We next investigated whether training separate single-task models for each kidney cell type provides any advantage over a single multi-task model. To do this, we trained ten independent models, one for each kidney cell type, using Enformer as the pretrained backbone because of its shorter input sequence length compared with Borzoi. We then compared the single-task and multi-task models across three evaluation tasks.

First, we assessed prediction accuracy on held-out genomic regions, including both ubiquitous and cell-type-specific peaks. For each model, predictions were generated separately for each cell type, and performance was measured using the Pearson correlation coefficient (*r*) between the observed and predicted chromatin accessibility. The two approaches achieved nearly identical performance in ubiquitous regions **(Fig. 6A)**. In cell-type-specific regions, single-task models scored slightly higher in most cell types, such as PT (*r* = 0.57 *vs*. 0.55), with larger gains in DT (*r* = 0.43 *vs*. 0.38) and particularly CD (*r* = 0.51 *vs*. 0.39) **(Fig. 6B)**. Next, we evaluated the ability of the models to predict allelic imbalance in DT, PT, and LOH cells. The performance of the single-task and multi-task models was almost identical. *AUROC* values were the same for PT and LOH (0.82) and differed only slightly for DT (0.83 for single-task *vs*. 0.82 for multi-task). Similarly, *AUPRC* values showed only marginal differences, with single-task models providing at most a slight improvement in some cases **(Fig. 6C,D)**. Finally, we evaluated both approaches for prioritizing GWAS fine-mapped causal variants in DT, PT, and LOH cells. Overall, the multi-task model showed slightly better performance, with only small differences compared to the single-task model. For instance, in DT and LOH cells, the multi-task model performed slightly better, achieving an *AUROC* of 0.78 and 0.79, respectively, compared to 0.76 and 0.78 for the single-task models. In PT cells, the multi-task model achieved an *AUPRC* of 0.46 compared to 0.45 for the single-task model **(Fig. 6E,F)**. Overall, single-task models did not provide a consistent performance advantage over the multi-task model for allelic-imbalance or GWAS variant prediction, and their gains in accessibility prediction were small, except in CD and DT.

**Figure 6:**
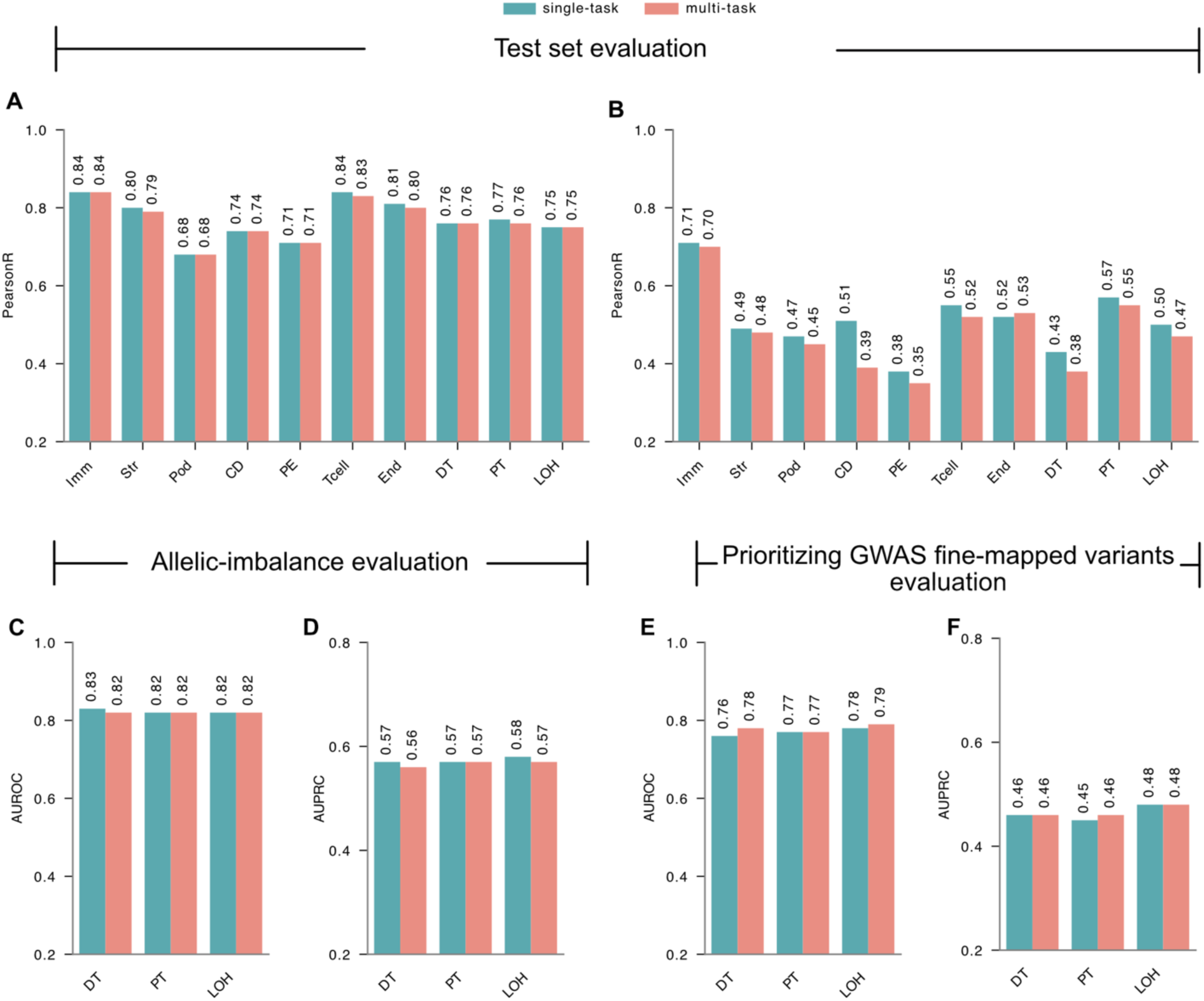
Comparison of single-task and multi-task models. **(A, B)** Test set evaluation comparing single-task and multi-task models across the ten kidney cell types. Performance is reported as the Pearson correlation (*r*) between observed and predicted chromatin accessibility for **(A)** ubiquitous genomic regions and **(B)** cell-type-specific regions. **(C, D)** Performance on allelic-imbalance prediction in DT, PT, and LOH cells, evaluated using **(C)** *AUROC* and **(D)** *AUPRC*. **(E, F)** Performance for prioritizing GWAS fine-mapped causal variants in DT, PT, and LOH cells, evaluated using **(E)** *AUROC* and **(F)** *AUPRC*.

## 3 Discussion

Training multi-task sequence models *de novo* is costly and design-intensive. Here we show that transfer learning from pretrained S2F models yields accurate kidney cell-type-specific models. While previous studies have reported limitations in predicting cell-type-specific accessibility with general-purpose models and in applying transfer learning to genomic tasks [45,46], we find that the representations learned by Borzoi and Enformer adapt effectively to kidney snATAC-seq, outperforming the specialized ChromKid model, especially in cell-type-specific chromatin accessibility prediction. These gains carry over to variant effect prediction. Kidzoi improves allelic-imbalance prediction over both general-purpose and specialized baselines in all three cell types tested, and Kidformer does so in PT and LOH.

S2F models have many applications, such as predicting variant effects, prioritizing fine-mapped causal variants, deciphering cis-regulatory logic, and designing functional DNA sequences. However, regardless of its benchmark score, a model that is excessively large or computationally slow has limited practical utility if it cannot be efficiently applied at scale. We show that 6–12% of the input (32–64*kbp* of local context) captures most of the information needed for variant effect prediction and for prioritization of GWAS fine-mapped causal variants, with diminishing returns beyond. These results suggest that locally focused models may offer a computationally cheaper alternative without meaningful loss of accuracy. Finally, while single-task models improved accessibility prediction, especially in CD and DT, they did not achieve a meaningful gain in allelic-imbalance or GWAS variant prediction despite far greater training complexity, favoring multi-task transfer as the more scalable strategy. Our results also demonstrate that the genomic aggregation window is an important consideration when applying large-context sequence models for variant prioritization. Although these models process hundreds of kilobases of genomic sequence, the choice of genomic region over which predictions are aggregated can substantially influence the ability to capture variant-specific regulatory effects.

To facilitate reuse by the community, we provide precomputed variant effect scores for ∼1.5 million common variants across all kidney cell types profiled in this study. We also release a PyTorch-native pipeline encompassing data preprocessing, model training, and variant scoring, designed to be readily extensible to other single-cell chromatin accessibility datasets.

Although transfer learning from pretrained models improves cell-type-specific model performance and variant effect prediction, a substantial performance gap remains between ubiquitous and cell-type-specific regulatory regions, underscoring the need for further methodological refinement to adequately capture context-specific regulatory grammar. Additionally, despite their predictive power, deep learning models remain largely difficult to interpret. Beyond these general challenges, our models are focused exclusively on chromatin accessibility, unlike multimodal models such as Borzoi and AlphaGenome. While they do not capture downstream molecular outcomes such as gene expression, focusing on a single modality can reduce complexity and allow more specialized modeling.

Future efforts could pair parameter-efficient fine-tuning with quantization and distillation to reduce model size and improve inference speed. Incorporating loss functions that explicitly reward correct rank-ordering of predictions across cell types, rather than focusing solely on absolute accuracy, may further improve both predictive performance and biological interpretability. Additionally, narrowing the gap between ubiquitous and cell-type-specific regions will likely depend on expanding training data for rare cell populations, or on developing model architectures that better represent combinatorial transcription factor activity. In summary, our work presents an accurate model for kidney tissue and shows that transfer learning is an effective and efficient tool for building cell-type-specific models from pretrained sequence models, and provides a resource and framework to extend this approach to other tissues and cell types.

## 4 Methods

### 4.1 Architectural details

#### 4.1.1 Borzoi

Borzoi is an S2F deep learning model that accepts genomic sequences of 524,288*bp* as input and predicts genome-wide coverage tracks at 32*bp* resolution. The architecture follows a U-Net-based encoder-decoder framework with integrated transformer layers, which progressively compresses the input sequence to 128*bp* for hierarchical motif extraction, captures long-range genomic interactions via self-attention [31], and reconstructs predictions at 32*bp* resolution through the decoder pathway. The final representations are projected through a 1 × 1 convolutional head to predict coverage tracks across hundreds of cell types outputting a 6,144 × 7,611 matrix. All Borzoi baseline scores were computed using the ensemble of four Borzoi replicates. We used the following script to score variants: https://github.com/calico/borzoi/blob/main/src/scripts/borzoi_sad.py.

#### 4.1.2 AlphaGenome

AlphaGenome is an S2F deep learning model that accepts genomic sequences of 1*Mbp* as input and predicts genome-wide coverage tracks at different resolutions ranging from 1*bp* to 2,048*bp*. The architecture follows a U-Net-based encoder-decoder framework with integrated transformer layers adapted from Borzoi, which progressively compresses the input sequence to 128*bp* for hierarchical motif extraction, captures long-range genomic interactions via self-attention, and reconstructs predictions at different resolutions through the decoder. The following PyTorch port was used to score variants: https://github.com/genomicsxai/alphagenome-pytorch, using the distilled all-fold checkpoint, a single model trained to approximate the predictions of the multi-fold ensemble.

#### 4.1.3 Enformer

Enformer is an S2F deep learning model that accepts genomic sequences of 196,608*bp* as input and predicts gene expression and chromatin accessibility profiles at 128*bp* resolution. The architecture consists of an initial convolutional stem followed by a series of residual convolutional blocks for local sequence feature extraction, succeeded by a stack of transformer attention blocks that capture long-range regulatory interactions across the input sequence. The final representations are projected through a pointwise linear head to predict coverage tracks across hundreds of cell types outputting an 896 × 5,313 matrix. We used the following script to score variants: https://github.com/calico/basenji/blob/master/bin/sonnet_sad.py.

#### 4.1.4 Sei

Sei is an S2F deep learning model that accepts genomic sequences of 4,096*bp* as input and predicts chromatin accessibility and regulatory activity profiles across 21,907 tracks. The architecture relies entirely on a hierarchical series of convolutional and dilated convolutional layers, which progressively expand the receptive field to capture both local sequence motifs and broader regulatory context without explicit attention mechanisms. The final representations are projected through a fully connected output head with a sigmoid activation function, producing predictions bounded between 0 and 1 so that each number is interpreted as an independent probability. We used the following script to score variants: https://github.com/FunctionLab/sei-framework/blob/main/1_variant_effect_prediction.py.

#### 4.1.5 ChromKid

ChromKid is a kidney-specific S2F deep learning model based on the Basset model [8] that accepts genomic sequences of 1,344*bp* as input and predicts chromatin accessibility across ten kidney cell types. The architecture consists of convolutional and dilated convolutional layers, progressively expanding the receptive field to capture local regulatory sequence features. The final representations are projected through an output head with a Softplus activation function, producing non-negative predictions for the central 192*bp* bin. We used the following script to score variants: https://github.com/ni-lab/kidney-finemapping/blob/main/kidney_finemapping/basenji/compute_sad.py.

### 4.2 Data processing

The data were processed using the same pipeline as the original Enformer and Borzoi studies. The input sequence lengths were unchanged: Kidzoi (based on Borzoi) used 524,288*bp* input sequences, while Kidformer (based on Enformer) used 196,608*bp* input sequences. The corresponding chromatin accessibility signals for training were prepared as follows. For Kidzoi, chromatin accessibility signals from a central region of 196,608*bp* of the input sequence, after cropping 163,840*bp* from each end, were aggregated at a 32*bp* resolution, yielding 6,144 output bins per track. For Kidformer, 40,960*bp* were cropped from each end, resulting in a central region of 114,688*bp* aggregated at a 128*bp* resolution, yielding 896 output bins per track. In both cases, the target chromatin accessibility signal from the pseudobulk bigWig files for the ten kidney cell types from Loeb et al. [30] was used for signal aggregation. We maintained the identical genomic region train/validation/test splits as used in the original Enformer and Borzoi studies, ensuring that no data leakage occurred between training and evaluation sets.

### 4.3 Model adaptation

We employed transfer learning from two pretrained genomic S2F models, Enformer and Borzoi, as backbone architectures using the PyTorch ports (Enformer: https://github.com/lucidrains/enformer-pytorch; Borzoi: https://github.com/johahi/borzoi-pytorch). We named the resulting transferred models Kidformer (Kidney-Enformer) and Kidzoi (Kidney-Borzoi), respectively. For adaptation to kidney chromatin accessibility data, the original task-specific prediction heads were removed and replaced with a randomly initialized output head mapping the backbone’s feature representations to *N* = 10 kidney cell type tracks. Specifically, a linear layer was used for Kidformer and a 1 × 1 convolutional layer for Kidzoi, consistent with the native architectural design of each backbone. Both were followed by a Softplus activation function to ensure non-negative predictions, appropriate for count-based chromatin accessibility data. All backbone weights were trainable during fine-tuning.

### 4.4 Model training

Genomic sequences were converted into a four-channel one-hot encoding. The bases A, C, G, and T correspond to [1,0,0,0], [0,1,0,0], [0,0,1,0], and [0,0,0,1], respectively. Unknown bases (N) were represented as [0,0,0,0]. The resulting input length was 196,608*bp* for fine-tuning Enformer and 524,288*bp* for fine-tuning Borzoi. To enhance model robustness and generalization, two data augmentation techniques were applied during the training phase. (1) Reverse-complement augmentation: With a 50% probability, the input sequence was subjected to reverse complementation, and the corresponding target profile was also reversed to maintain strand consistency. (2) Shift augmentation: A random uniform shift of ±3*bp* was applied to the sequence window coordinates for each training example.

The training objective was defined by the Poisson negative log-likelihood loss function. We utilized the fused AdamW optimizer algorithm [47] with standard hyperparameters *β*_1_ = 0.9, *β*_2_ = 0.999, and *ε* = 1 × 10^−8^. To implement differential weight decay, model parameters were grouped by dimensionality: all parameters with two or more dimensions were assigned a weight decay of 0.1, while parameters with fewer than two dimensions were assigned a weight decay of 0.0. The learning rate schedule included a linear warmup phase, where the rate increased from zero to a peak value of 3 × 10^−4^ over the first 1,000 optimization steps, followed by a cosine decay to a minimum floor of 3 × 10^−5^ over the remaining steps. Training was performed on a single NVIDIA A100 GPU with a per-device batch size of 8 and 2 for Kidformer and Kidzoi, respectively, with an effective batch size of 64 achieved through gradient accumulation. Gradient clipping with a maximum L_2_-norm of 0.2 was applied to ensure numerical stability. Mixed-precision training using the *bfloat16* data type was employed to accelerate computation and reduce memory overhead. Models were trained for 8 epochs, and the checkpoint with the lowest validation loss was retained. Four replicate models of Kidzoi were trained, each initialized from the corresponding Borzoi replicate. Unless stated otherwise, all analyses in this study used replicate rep0, and the released weights on Zenodo include all four replicates, the single Kidformer model, and the ten single-task models. The ten single-task models used in the multi-task comparison were trained with the same hyperparameters, schedule and data splits as Kidformer, each with a one-track output head for its cell type.

### 4.5 Cell-type-specific and ubiquitous evaluation

The test set sequences were stratified into two distinct groups: regions exhibiting activity ubiquitously across all ten cell types and those classified as cell-type-specific from Loeb et al. [30].

Performance for the Kidzoi and Kidformer models was evaluated separately on these two sets. To obtain predictions, ubiquitous peaks were processed using a single forward pass, whereas cell-type-specific peaks required a separate forward pass for each cell type as the active regions differ between cell types. Following prediction, the specific targets were extracted and aligned with the corresponding center predictions to compute the Pearson correlation coefficient. This calculation was strictly limited to a consistently sized 256*bp* central region of the model output to standardize comparison across differing output resolutions. For the Kidformer model (896 total bins), this evaluation region comprised the central two bins, while for the Kidzoi model (6,144 total bins) it corresponded to the central eight bins. Only test-set regions with an observed signal greater than 1 were included, and Pearson *r* was computed between log(*x* + 1)-transformed predicted and observed signals.

### 4.6 Allelic imbalance evaluation

Allelic imbalance data were obtained from Loeb et al. [30]. Briefly, the authors processed variant data to define high-confidence positive (imbalanced) and negative (balanced) sets for PT, LOH, and DT. Raw allelic imbalance variants were first filtered to retain only biallelic SNVs with a total read depth greater than 20. Indels and triallelic sites were excluded. To ensure functional relevance, analysis was restricted to variants intersecting with open chromatin peaks. Significant allelic imbalance was identified by correcting *P*-values for multiple testing using the Benjamini–Hochberg procedure. Variants exhibiting a false discovery rate (FDR) < 0.01 were classified as the positive set. A matched negative control set was defined using the remaining variants with *P* > 0.1. To control for coverage bias, these balanced sites were stratified into 20 bins based on log-transformed total read counts. Within each bin, negative variants were randomly sampled to mirror the coverage distribution of the positive set, yielding a final negative set seven times larger than the positive set.

For each variant, reference and alternative sequences were extracted, centered on the variant position using input windows of 196,608*bp* and 524,288*bp* for Kidformer and Kidzoi, respectively. To reduce positional sensitivity, shift augmentation of [−1, 0, +1] *bp* was applied, and predictions were averaged across all shifts. To ensure strand invariance, predictions were additionally generated for both forward and reverse-complement orientations and averaged across strands for each allele. For each allele, the predicted signal was aggregated over a 256*bp* window centered on the variant (eight Kidzoi bins and two Kidformer bins), giving ref and alt scores. Allelic imbalance was then calculated as: 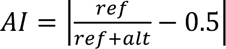. For the aggregation-window analysis, the output cropping was removed so that predictions spanned the full input sequence (16,384 bins for Kidzoi and 1,536 bins for Kidformer); “full” refers to this entire span. To evaluate effect direction, variants were labeled positive when the observed reference-allele fraction, ref/(ref + alt), exceeded 0.5 and negative otherwise, and were scored by the predicted reference-allele fraction.

### 4.7 GWAS fine-mapped variant evaluation

Functionally informed fine-mapped variants derived from an eGFRcr-cys GWAS were obtained from Loeb et al. [30]. Variant coordinates were lifted over from hg19 to hg38 using CrossMap [48], and coding variants and indels were excluded, retaining only non-coding fine-mapped SNVs with posterior inclusion probability (PIP) > 0.3 as the positive set. A negative set of MAF-matched SNVs, seven times the size of the positive set, was sampled from the full GWAS summary statistics, excluding variants overlapping any fine-mapped locus and restricting to those with *P* > 0.5. This negative set was generated independently as part of the present study and was not provided by Loeb et al. [30]. Similar to the allelic imbalance evaluation, reference and alternative sequences were extracted, centered on the variant position using input windows of 196,608*bp* and 524,288*bp* for Kidformer and Kidzoi, respectively. Shifts of [−1, 0, +1] *bp* were applied, and predictions were averaged across all shifts and over forward and reverse-complement orientations. For each variant, the SNP activity difference (SAD) score was computed as the difference between the summed alternative and reference predicted chromatin accessibility signals, aggregated from the central 1*kbp* of the model output. Variants were ranked by the absolute value of the SAD score.

### 4.8 Precomputed variant scores

We downloaded the full set of common variants from Srivastava et al. [49] and performed an intersection with the open chromatin regions identified in the snATAC-seq data from Loeb et al. [30], yielding ∼1.5 million variants. We then scored these variants with Kidzoi in all ten kidney cell types, using the SAD procedure described above over a central 1*kbp* region and averaging predictions across the forward and reverse-complement strands. The scores are available on Zenodo (see Data and code availability).

### 4.9 Relative performance gains

Relative gains were calculated from the reported values as 100 × (x − y)/(y − b), where x is the score being compared (for example, Kidzoi, or a model at a longer context), y is the reference score (a baseline model, or a shorter context), and b is the score of a random predictor (0 for Pearson *r*, 0.5 for *AUROC*, and the positive-class prevalence, 0.125, for *AUPRC*), and were rounded to the nearest whole percent.

## Acknowledgments

We acknowledge the Boston Children’s Hospital High-Performance Computing (BCH HPC) resources, including the E3 cluster and associated GPU computing resources, for supporting the computational analyses performed in this study. We also thank the members of the Sampson and Lee labs for their valuable discussions, feedback, and support throughout the course of this work. We thank Gabriel Loeb for his valuable comments and feedback.

## Author contributions

Conceptualization: AAR, DL

Data curation: AAR

Formal analysis: AAR

Methodology: AAR, DL

Investigation: AAR

Visualization: AAR

Funding acquisition: DL

Project administration: AAR, DL

Resources: AAR

Software: AAR

Validation: AAR

Supervision: DL

Writing – original draft: AAR, DL

Writing – review & editing: AAR, DL

## Data and code availability

All the data to train and evaluate models were obtained from Loeb et al. [30], including the snATAC-seq data (GEO accession GSE262931) and the eGFRcr-cys GWAS summary statistics (GWAS Catalog accession GCST90428446). The model weights, scores, and evaluation results are available on Zenodo: https://doi.org/10.5281/zenodo.22391344. The code for training and using the models, as well as the code used to generate figures, is available at https://github.com/dongwonlee-lab/kidzoi-kidformer.

## Writing

A large language model (LLM) was used to assist with improving the clarity and phrasing of the manuscript. All LLM-generated suggestions were critically reviewed, revised, and approved by the authors.

## Ethics statement

This study analyzed previously published, de-identified human kidney single-nucleus ATAC-seq and GWAS summary data. No new human-subject data were generated, and no additional ethical approval was required.

