## Supplemental File for "Kidzoi Enables Cell-Type-Specific Regulatory Activity and Variant Effect Prediction in Kidney"

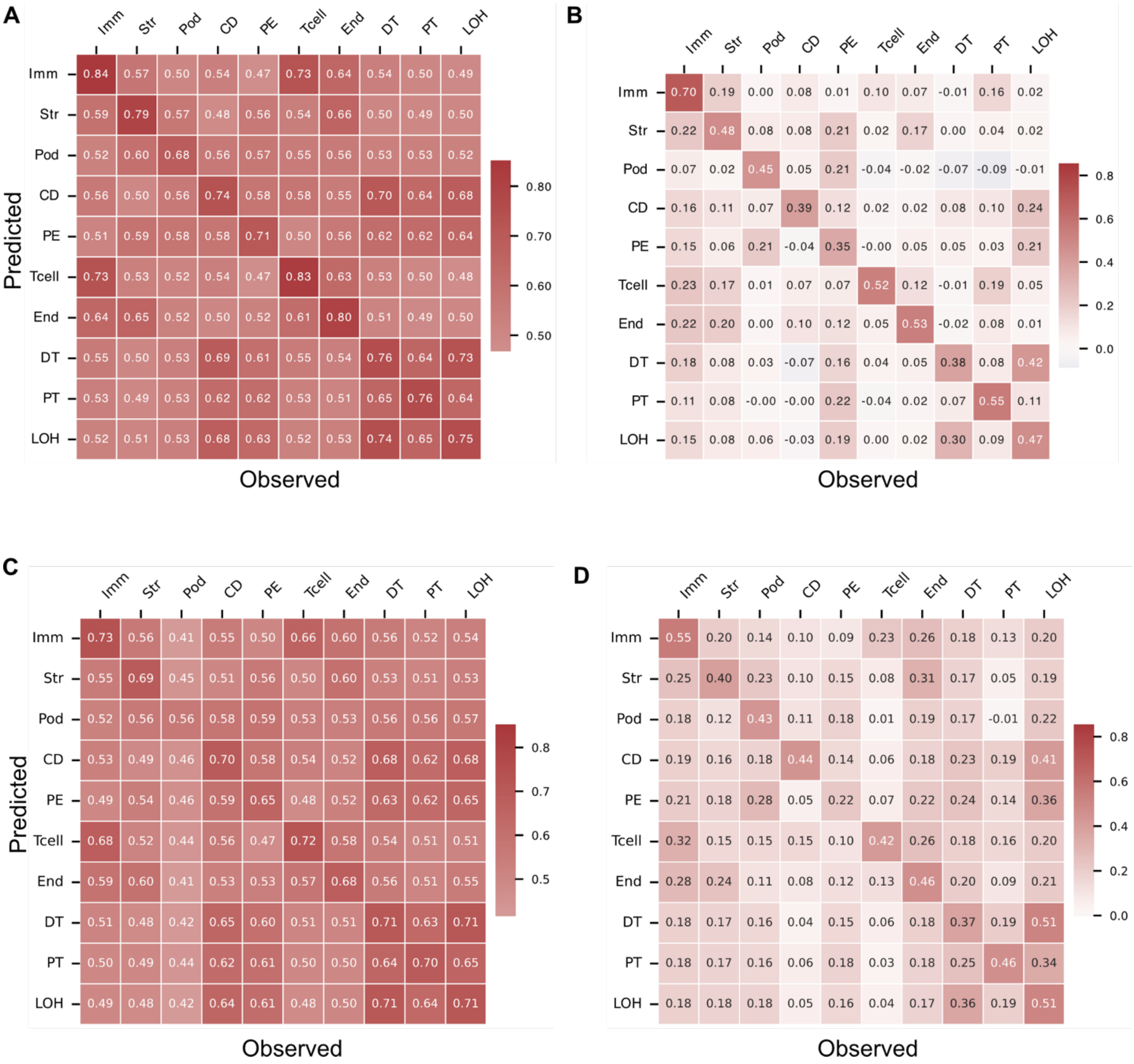


**Figure S1: Performance of Kidformer and ChromKid in predicting ubiquitous and cell-type-specific regions. (A, B)** Heatmaps of Kidformer performance (Pearson correlation) in ubiquitous (A) and cell-type-specific (B) regions across all cell types. Each entry $(i, j)$ represents the Pearson $r$ between predicted signal in cell type $i$ and the observed signal for cell type $j$; diagonal entries are within-cell-type, off-diagonal entries cross-cell-type. **(C, D)** Heatmaps of ChromKid in ubiquitous (C) and cell-type-specific (D) regions. Cell-type abbreviations: Imm, immune; Str, stromal; Pod, podocyte; CD, collecting duct; PE, parietal epithelial; Tcell, T cell; End, endothelial; DT, distal tubule; PT, proximal tubule; LOH, loop of Henle. The same abbreviations are used throughout the Supplementary Material.

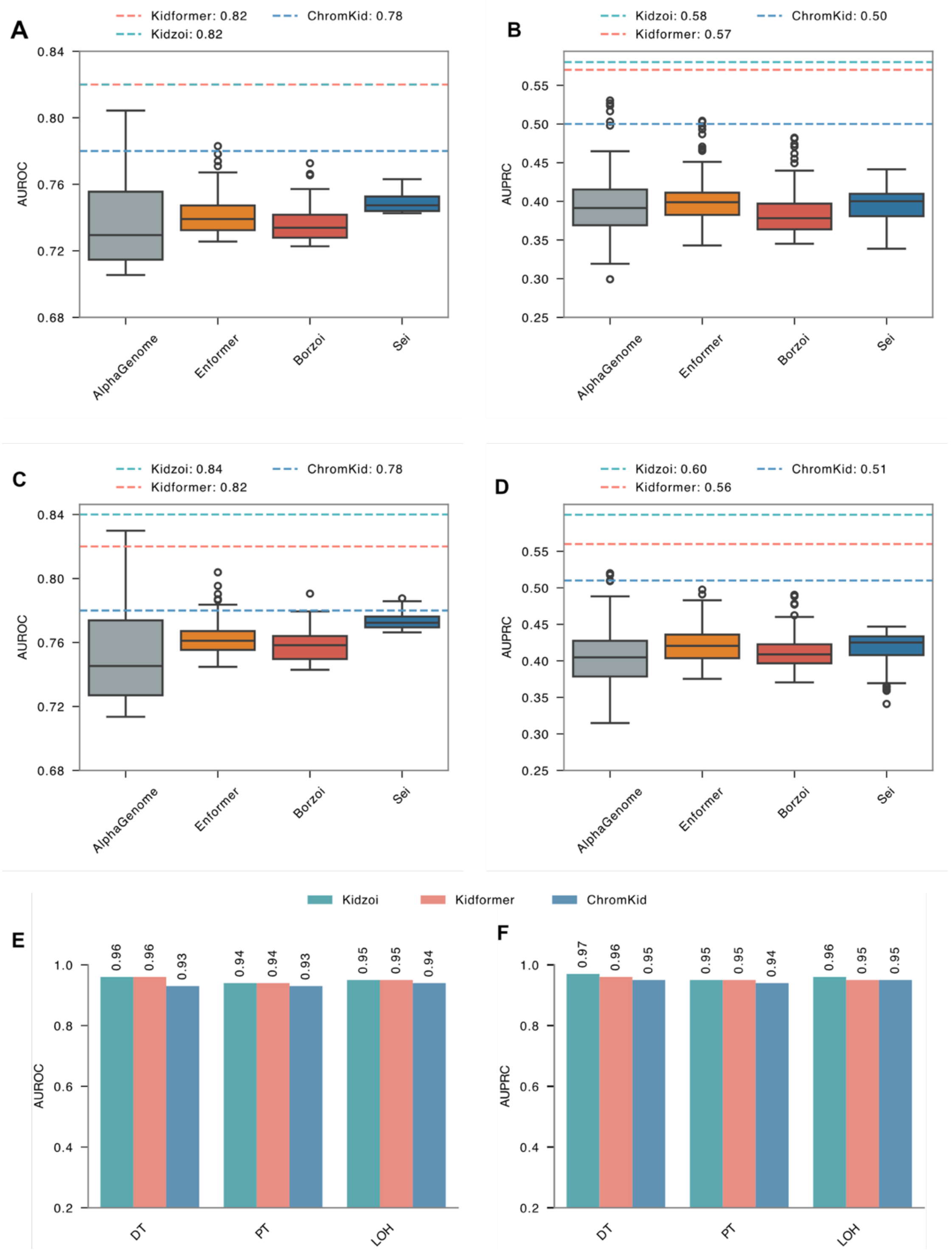


**Figure S2: Performance of pretrained and transferred models in predicting allelic imbalance and directionality**. **(A–D)** Box plots showing performance distributions (*AUROC* and *AUPRC*) across the top 100 DNase and ATAC tracks from each pretrained model, while dashed horizontal lines indicate the performance of transferred and specialized models as single reference values in LOH (**A, B**) and DT **(C, D)**. **(E, F)** Bar plots showing how accurately Kidzoi, Kidformer, and ChromKid can predict directionality across all three cell types in terms of *AUROC* (E) and *AUPRC* (F).

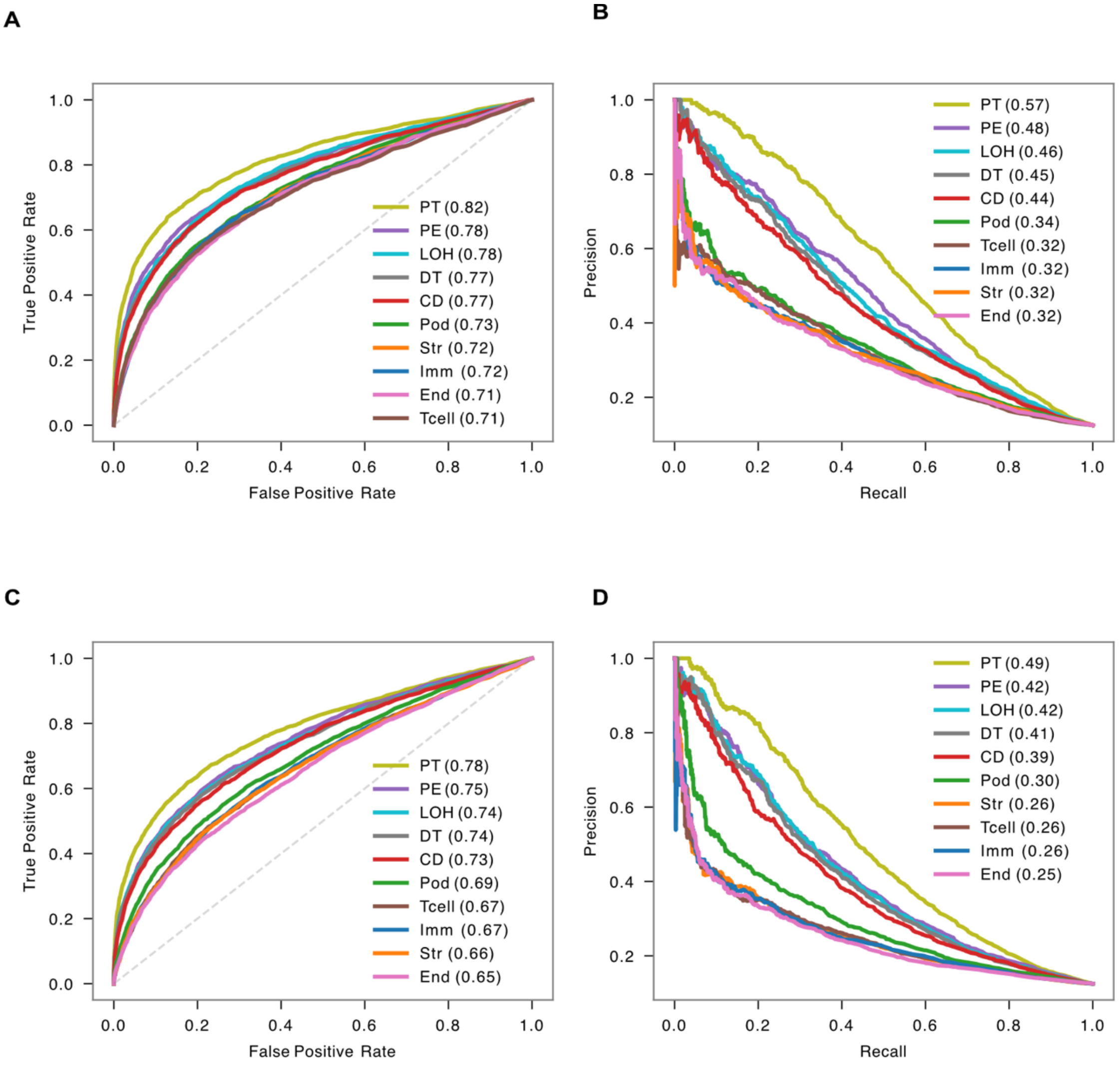


**Figure S3: Performance of Kidformer and ChromKid in predicting variants with allelic imbalance in proximal tubule.** ROC and precision-recall curves of Kidformer **(A, B)** and ChromKid **(C, D)** for the ten kidney cell types.

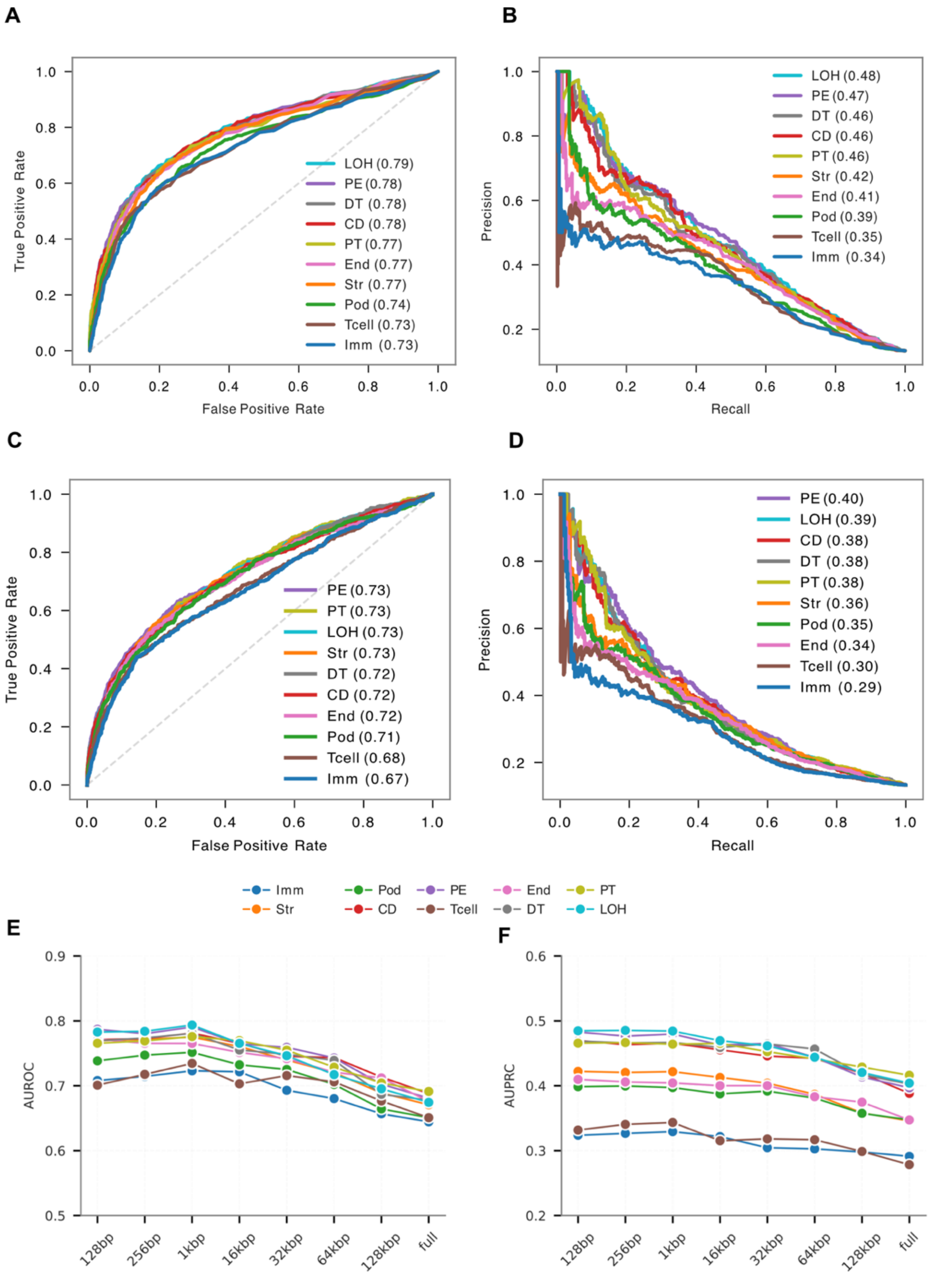


**Figure S4: Performance of Kidformer and ChromKid in predicting GWAS fine-mapped variants.** ROC and precision-recall curves of Kidformer **(A, B)** and ChromKid **(C, D)** for the ten kidney cell types. (**E, F)** Line plots of Kidformer performance as a function of signal aggregation window size around the variant position across ten cell types. *AUROC* peaks at or near 1*kbp* and declines at larger windows, whereas *AUPRC* is relatively stable from 128*bp* to 1*kbp* and generally decreases at larger windows.

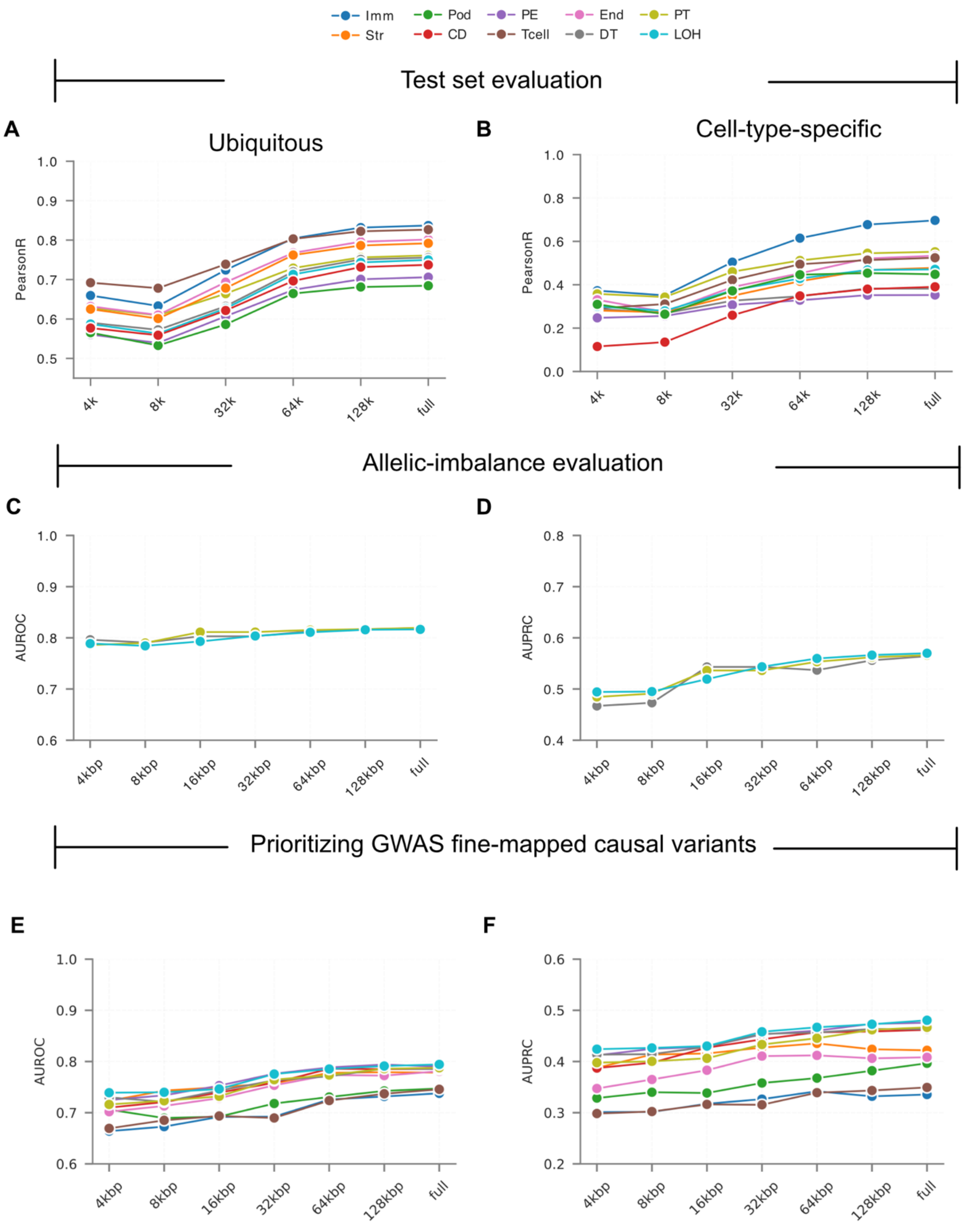


**Figure S5: The effect of input sequence truncation on Kidformer performance**. Performance (Pearson correlation) in predicting **(A)** ubiquitous regions and **(B)** cell-type-specific regions. **(C)** *AUROC* and **(D)** *AUPRC* for predicting allelic imbalance. **(E)** *AUROC* and **(F)** *AUPRC* for predicting GWAS fine-mapped variants. The x-axis shows the length of the actual genomic sequence around the variant, with the remaining positions padded with *Ns* to maintain the full 196,608*bp* input length.

**Table S1:** Test-set Pearson correlation for ubiquitous open chromatin region prediction across different sequence context lengths using Kidzoi.

| Context | Imm | Str | Pod | CD | PE | Tcell | End | DT | PT | LOH |
|---|---|---|---|---|---|---|---|---|---|---|
| 4*kbp* | 0.636 | 0.591 | 0.554 | 0.633 | 0.483 | 0.647 | 0.603 | 0.633 | 0.614 | 0.607 |
| 8*kbp* | 0.662 | 0.616 | 0.560 | 0.638 | 0.501 | 0.681 | 0.627 | 0.642 | 0.619 | 0.609 |
| 16*kbp* | 0.703 | 0.659 | 0.578 | 0.664 | 0.535 | 0.715 | 0.657 | 0.671 | 0.641 | 0.638 |
| 32*kbp* | 0.753 | 0.697 | 0.603 | 0.704 | 0.585 | 0.755 | 0.698 | 0.712 | 0.675 | 0.681 |
| 64*kbp* | 0.790 | 0.724 | 0.624 | 0.726 | 0.633 | 0.784 | 0.731 | 0.738 | 0.708 | 0.712 |
| 128*kbp* | 0.817 | 0.752 | 0.641 | 0.743 | 0.664 | 0.803 | 0.757 | 0.753 | 0.737 | 0.730 |
| 256*kbp* | 0.832 | 0.772 | 0.654 | 0.756 | 0.691 | 0.818 | 0.778 | 0.766 | 0.760 | 0.751 |
| 524*kbp* | 0.837 | 0.782 | 0.665 | 0.764 | 0.702 | 0.824 | 0.786 | 0.774 | 0.770 | 0.761 |

**Table S2:** Test-set Pearson correlation for cell-type-specific open chromatin region prediction across different sequence context lengths using Kidzoi.

| Context | Imm | Str | Pod | CD | PE | Tcell | End | DT | PT | LOH |
|---|---|---|---|---|---|---|---|---|---|---|
| 4*kbp* | 0.305 | 0.323 | 0.366 | 0.441 | 0.144 | 0.115 | 0.327 | 0.357 | 0.409 | 0.419 |
| 8*kbp* | 0.314 | 0.339 | 0.364 | 0.473 | 0.159 | 0.161 | 0.348 | 0.362 | 0.408 | 0.422 |
| 16*kbp* | 0.381 | 0.376 | 0.390 | 0.538 | 0.188 | 0.255 | 0.377 | 0.380 | 0.421 | 0.446 |
| 32*kbp* | 0.467 | 0.412 | 0.427 | 0.587 | 0.222 | 0.382 | 0.415 | 0.401 | 0.459 | 0.491 |
| 64*kbp* | 0.552 | 0.452 | 0.456 | 0.607 | 0.276 | 0.474 | 0.453 | 0.422 | 0.511 | 0.519 |
| 128*kbp* | 0.618 | 0.490 | 0.479 | 0.615 | 0.326 | 0.535 | 0.506 | 0.426 | 0.557 | 0.529 |
| 256*kbp* | 0.660 | 0.512 | 0.498 | 0.621 | 0.361 | 0.574 | 0.541 | 0.428 | 0.593 | 0.542 |
| 524*kbp* | 0.677 | 0.529 | 0.508 | 0.624 | 0.368 | 0.592 | 0.560 | 0.434 | 0.610 | 0.557 |

**Table S3:** *AUROC* performance for allelic imbalance prediction in DT, PT, and LOH kidney cell types across different sequence context lengths using Kidzoi.

| Context | DT | PT | LOH |
|---|---|---|---|
| 4*kbp* | 0.743 | 0.703 | 0.749 |
| 8*kbp* | 0.764 | 0.734 | 0.771 |
| 16*kbp* | 0.784 | 0.767 | 0.788 |
| 32*kbp* | 0.795 | 0.789 | 0.807 |
| 64*kbp* | 0.806 | 0.805 | 0.810 |
| 128*kbp* | 0.809 | 0.810 | 0.813 |
| 256*kbp* | 0.811 | 0.815 | 0.814 |
| 524*kbp* | 0.812 | 0.816 | 0.814 |

**Table S4:** *AUPRC* performance for allelic imbalance prediction in DT, PT, and LOH kidney cell types across different sequence context lengths using Kidzoi.

| Context | DT | PT | LOH |
|---|---|---|---|
| 4*kbp* | 0.432 | 0.300 | 0.431 |
| 8*kbp* | 0.461 | 0.384 | 0.460 |
| 16*kbp* | 0.508 | 0.445 | 0.500 |
| 32*kbp* | 0.531 | 0.512 | 0.536 |
| 64*kbp* | 0.549 | 0.527 | 0.552 |
| 128*kbp* | 0.562 | 0.543 | 0.558 |
| 256*kbp* | 0.565 | 0.554 | 0.563 |
| 524*kbp* | 0.562 | 0.557 | 0.563 |

**Table S5:** *AUROC* performance for GWAS fine-mapped variant prediction across kidney cell types using Kidzoi model across different sequence context lengths.

| Context | Imm | Str | Pod | CD | PE | Tcell | End | DT | PT | LOH |
|---|---|---|---|---|---|---|---|---|---|---|
| 4*kbp* | 0.588 | 0.675 | 0.658 | 0.663 | 0.705 | 0.596 | 0.621 | 0.699 | 0.671 | 0.686 |
| 8*kbp* | 0.614 | 0.695 | 0.661 | 0.689 | 0.717 | 0.625 | 0.647 | 0.722 | 0.692 | 0.710 |
| 16*kbp* | 0.643 | 0.711 | 0.701 | 0.717 | 0.735 | 0.654 | 0.672 | 0.736 | 0.703 | 0.739 |
| 32*kbp* | 0.676 | 0.731 | 0.731 | 0.749 | 0.762 | 0.667 | 0.704 | 0.755 | 0.734 | 0.756 |
| 64*kbp* | 0.694 | 0.762 | 0.737 | 0.763 | 0.771 | 0.688 | 0.726 | 0.761 | 0.749 | 0.759 |
| 128*kbp* | 0.699 | 0.768 | 0.745 | 0.765 | 0.778 | 0.704 | 0.746 | 0.767 | 0.764 | 0.772 |
| 256*kbp* | 0.710 | 0.771 | 0.739 | 0.768 | 0.781 | 0.710 | 0.753 | 0.771 | 0.777 | 0.783 |
| 524*kbp* | 0.709 | 0.770 | 0.747 | 0.766 | 0.777 | 0.711 | 0.755 | 0.776 | 0.777 | 0.783 |

**Table S6:** *AUPRC* performance for GWAS fine-mapped variant prediction across kidney cell types using Kidzoi model across different sequence context lengths.

| Context | Imm | Str | Pod | CD | PE | Tcell | End | DT | PT | LOH |
|---|---|---|---|---|---|---|---|---|---|---|
| 4*kbp* | 0.259 | 0.373 | 0.330 | 0.371 | 0.401 | 0.266 | 0.292 | 0.409 | 0.372 | 0.412 |
| 8*kbp* | 0.279 | 0.387 | 0.335 | 0.396 | 0.417 | 0.296 | 0.317 | 0.430 | 0.397 | 0.429 |
| 16*kbp* | 0.309 | 0.402 | 0.365 | 0.417 | 0.424 | 0.319 | 0.351 | 0.436 | 0.411 | 0.448 |
| 32*kbp* | 0.332 | 0.409 | 0.376 | 0.438 | 0.445 | 0.331 | 0.369 | 0.451 | 0.442 | 0.464 |
| 64*kbp* | 0.337 | 0.417 | 0.374 | 0.439 | 0.448 | 0.337 | 0.379 | 0.446 | 0.446 | 0.460 |
| 128*kbp* | 0.329 | 0.411 | 0.374 | 0.430 | 0.448 | 0.342 | 0.382 | 0.445 | 0.447 | 0.458 |
| 256*kbp* | 0.328 | 0.409 | 0.369 | 0.429 | 0.452 | 0.346 | 0.385 | 0.446 | 0.448 | 0.460 |
| 524*kbp* | 0.331 | 0.406 | 0.371 | 0.431 | 0.448 | 0.344 | 0.384 | 0.447 | 0.445 | 0.460 |
